# Telomerase inhibition causes premature senescence of fetal guinea pig muscle cells

**DOI:** 10.1101/2020.02.21.959320

**Authors:** Stephanie E. Hallows, Timothy R.H. Regnault, Dean H. Betts

## Abstract

Premature senescence in low birth weight rodents is associated with later life metabolic disease, including the development of insulin resistance. Telomerase, a reverse transcriptase enzyme with telomeric and non-telomeric functions, is present at high levels during development to maintain and repair long telomere lengths and to protect cells from oxidative stress-induced growth arrest. Adverse *In utero* environments are often associated with increased reactive oxygen species (ROS), and ROS have been documented to impair/alter telomerase function. We postulate that telomerase protects cells against oxidative stress-induced damage, and its inhibition could lead to premature senescence. A primary cell line of fetal guinea pig muscle cells was differentiated under high (20%) and low (1-2%) oxygen concentrations and telomerase activity was pharmacologically inhibited using a synthetic tea catechin. Following 48 hours, ROS detection was conducted with MitoSOX, Mitotracker and 6-carboxy-2’,7’-dichlorodihydrofluorescein diacetate staining. Cells cultured at 20% O_2_ and treated with a telomerase activity inhibitor displayed reduced cell growth rates and increased levels of senescence markers, including p21 and p53. Telomeric DNA damage, measured by phosphorylated-γH2A.X staining at telomeres, was significantly increased in cells cultured at all oxygen concentrations with telomerase inhibition. Telomerase inhibition altered metabolic signaling (e.g. mTOR, p66Shc) and increased mitochondrial ROS levels. Telomerase may protect cells during development from adverse *in utero* environments that cause premature senescence.

## Introduction

A great number of human epidemiological and animal model studies have reported strong associations between the birth phenotype, particularly birth weight, and risk for many complex common adult diseases in later life, including hypertension, coronary artery disease and diabetes [1, 2]. Adverse intrauterine environments due to maternal malnutrition, hypertension, and placental insufficiency cause alterations in oxygen and nutrient supply that results in intrauterine growth restriction (IUGR) where a fetus fails to reach its genetic growth potential [3–5]. The redistribution of nutritional resources leads to a decrease in muscularity and an increase in the percentage of body fat in these infants that lasts throughout childhood and adult life [6, 7] in association with changes in insulin sensitivity and other markers of metabolic syndrome [8, 9]. The effects of impaired growth during *in utero* life are exacerbated by a rapid postnatal growth [10, 11] and appear to predispose offspring to an increased risk of disease in later life, or the concept of developmental origins of health and disease (DOHaD)[12].

Although the molecular and cellular mechanisms underlying the association between birth size and early onset of age-related disease are currently unclear, a number of studies indicate that reprogramming of the hypothalamic-pituitary-adrenal axis [13, 14], altered insulin signaling pathways [15, 16] and epigenetic modifications [17, 18] are involved. Emerging data suggest that oxidative stress and mitochondrial dysfunction may also play a major role in adverse IUGR outcomes [19, 20]. Although there are most likely separate disease-specific processes at play, there may be common underlying mechanisms mediating the effects of a diverse intrauterine perturbations on a range of health and disease risk outcomes.

One stress sensor that links the surrounding microenvironment to cellular function/state is the telomere. Mammalian telomeres contain a six base-pair sequence (TTAGGG)^n^ that is tandemly repeated many kilobases at both ends of every chromosome [21, 22]. Telomeres are essential in maintaining chromosome stability, playing a vital role in the correct segregation of chromosomes during cell division [23, 24] and counteracting the loss of terminal DNA sequences that occurs during DNA synthesis [25]. Telomere dysfunction that has been linked to cellular aging by acting as a “mitotic clock” causes cells to enter a permanent cell growth arrest state (senescence) when a “critically” short telomere length is reached [26] or when the telomere capping structure has been disrupted by triggers such as oxidative stress [27–29].

Telomere shortening/dysfunction can be overcome by the expression of the ribonucleoprotein enzyme telomerase [30]. This multi-subunit reverse transcriptase uses its RNA component (TR or TERC) to align the telomerase reverse transcriptase (TERT) component to the chromosomal ends and acts as a template for the addition of telomeric DNA [31, 32]. Telomerase activity has been detected in germ cells, cells of renewal tissues, and cancer cells but not in most normal somatic tissues [33–35]. Telomerase provides an extended replicative life span while preserving telomere integrity [36–38], while under oxidative stress conditions telomerase is shuttled out of the nucleus to protect mitochondrial function to reduce ROS-induced telomeric and non-telomeric DNA damage at the cost of telomere shortening [39, 40].

Shortened telomeres, and/or reduced telomerase production have been linked to shortened life span [41–43] and several age-related diseases, including hypertension, cardiovascular disease, and type 2 diabetes [44–47]. Recently, telomere length/integrity has emerged beyond its accepted role as a biomarker of cellular aging and senescence to one that appears to play a more causative role in cellular regeneration, metabolism, aging and disease [48, 49]. Recent studies suggest that telomeric integrity not only affects cellular replicative capacity but also underlies the execution of global epigenetic changes that affects cell signaling, cellular senescence and aging [50–52]. Thus, alterations in the initial setting of telomere length and/or production of telomerase could impact postnatal health and longevity.

Human and animal studies have found an association between adverse or suboptimal intrauterine environments and shorter offspring telomere length [53–55]. Stressors including hypoxia and altered nutrient availability encountered *in utero* increase oxidative stress levels in low birth weight individuals [56–58] with elevated senescent marker expression in various postnatal tissues [55, 59–61]. In addition, telomerase subunits and activity have been found decreased in IUGR placentas [59, 62, 63]. Telomerase may play a role in protecting cells from cellular senescence when exposed to stressful conditions through its telomeric and extra-telomeric roles in DNA damage response, apoptosis protection and modulation of oxidative stress and mitochondrial function [64, 65]. The aim of this study was to modulate telomerase activity levels in fetal muscle cells cultured under varying oxidative stress conditions as an *in vitro* proxy to determine the downstream effects of telomerase inhibition on cellular senescence, DNA damage, intracellular ROS levels and metabolism pathways in a cell type that is affected by IUGR.

## Results

### Confirmation of fetal myoblast lineage

Muscle transcription factor markers Pax7, myogenin and MyoD were used to confirm the identity of the primary fetal guinea pig myoblast cell line. These transcription factors are expressed at different points during myoblast differentiation, and can be used to determine stage of muscle differentiation [66]. Fetal guinea pig muscle cells were cultured at 20% O_2_ for 48 hrs before fixing and staining with lineage-specific antibodies. Fluorescence microscopy images showed nuclear localization of each muscle-specific transcription factor, indicating a myoblast cell lineage (S1 Fig). Pax7 regulates the transcription of differentiation factors in muscle progenitor cells, and showed the weakest staining. MyoD and myogenin are necessary for myogenic determination at the myoblast stage, and were both highly expressed and localized to the nucleus in fetal guinea pig muscle cells.

### Low oxygen culture is refractory to pharmacological telomerase activity inhibition

Fetal guinea pig muscle cells were cultured at both high (20%) O_2_ or low (2%) O_2_ tensions and were treated with increasing dosages of Telomerase Inhibitor (TI)-IX from 0.1 μM to 2.0 μM for 48 hrs (Fig 1). At 20% O_2_ tensions, telomerase activity was significantly decreased in fetal guinea pig muscle cells with 0.5 μM and 1.0 μM TI-IX (p<0.05) and with 2.0 μm TI-IX (p<0.01) treatments. In fetal guinea pig muscle cells cultured at 2% O_2_ tensions, telomerase activity was significantly decreased with 2.0 μM treatment of TI-IX (p<0.05), but had significantly (p<0.05) higher telomerase activity levels than 20% O_2_ cultured cells exposed to lower TI-IX concentrations. This significant difference in telomerase activity levels between oxygen concentrations was apparent at the different Telomerase Inhibitor IX treatments (p<0.05) except at the highest dose of 2.0μM (Fig 1).

**Fig 1.**
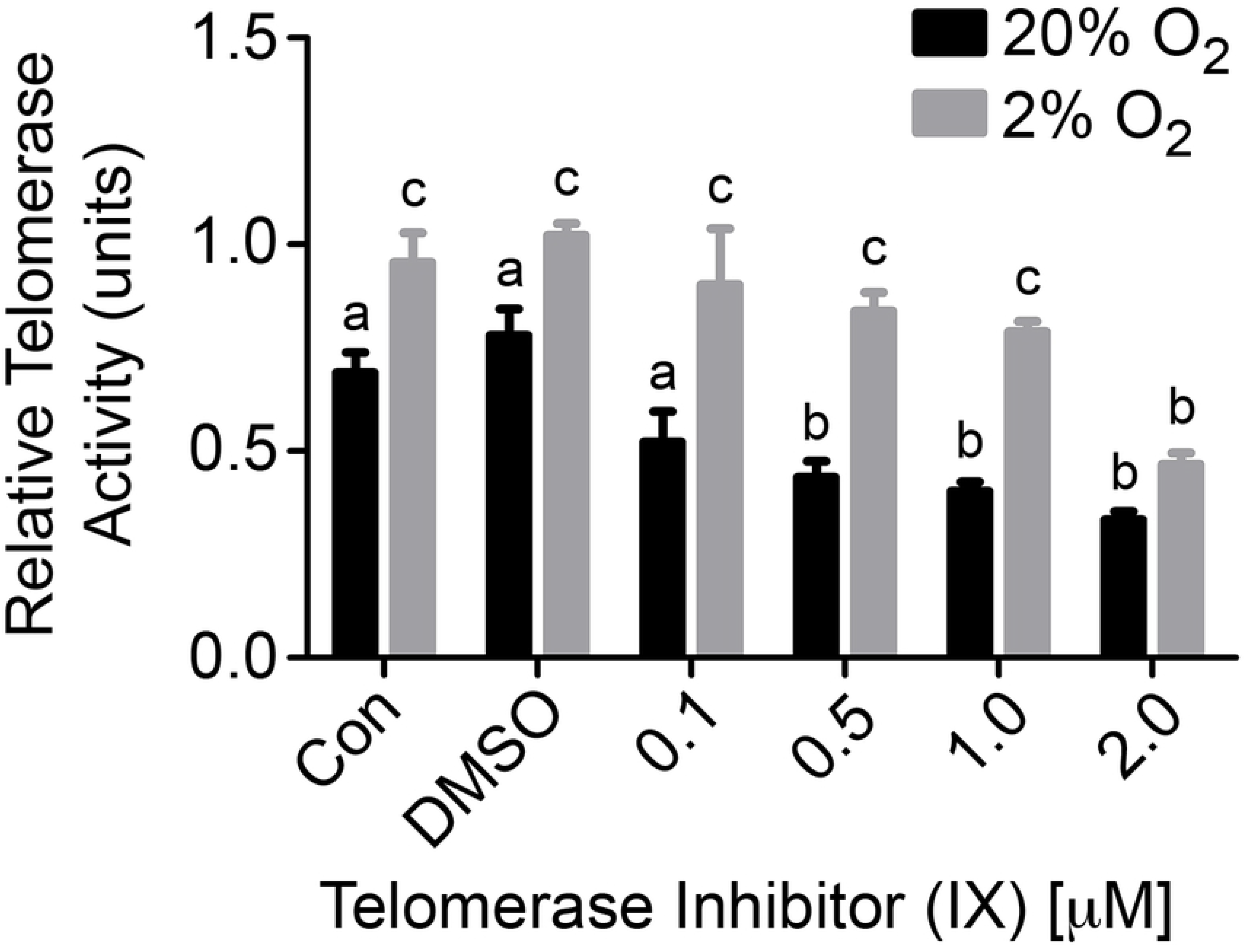
Low oxygen tension is refractory to pharmacological telomerase inhibition in fetal guinea pig muscle cells. Fetal guinea pig muscle cells were treated with increasing concentrations of a synthetic tea catechin (telomerase inhibitor (TI)-IX; Calbiochem) for 48 hours of culture under 2% and 20% oxygen (O_2_) tensions. Telomerase activity was measured using the Real-time Quantitative Telomeric Repeat Amplification Protocol (RQ-TRAP) assay. Telomerase activity levels significantly decreased with 0.5µM, 1.0µM and 2.0µM TI-IX treatments compared to non-treated (Con) and vehicle-treated (DMSO) cells cultured in 20% O_2_ conditions. Telomerase activity was significantly decreased with 2.0µM TI-IX compared to controls. Cells cultured in 2% O_2_ conditions exhibited significantly higher telomerase activity levels in control cell groups and cells treated with TI-IX at every concentration, except for 2.0µM, compared to 20% O_2_ grown/treated cells. Results (n=3) were analyzed with a two-way ANOVA and different letters above the histogram bars represent significant differences (p<0.05) in mean telomerase activities.

### Senescent characteristics displayed by 20% oxygen cultured cells treated with telomerase inhibitor

Fetal guinea pig muscle cells treated with increasing concentrations of telomerase inhibitor IX (TI-IX) displayed altered cell morphology including larger, flatter cells and reduced confluency that are typical morphological characteristics of senescent cells (Fig 2). This effect was observed to a greater extent at 20% O_2_ compared to 2% O_2_ culture conditions.

**Fig 2.**
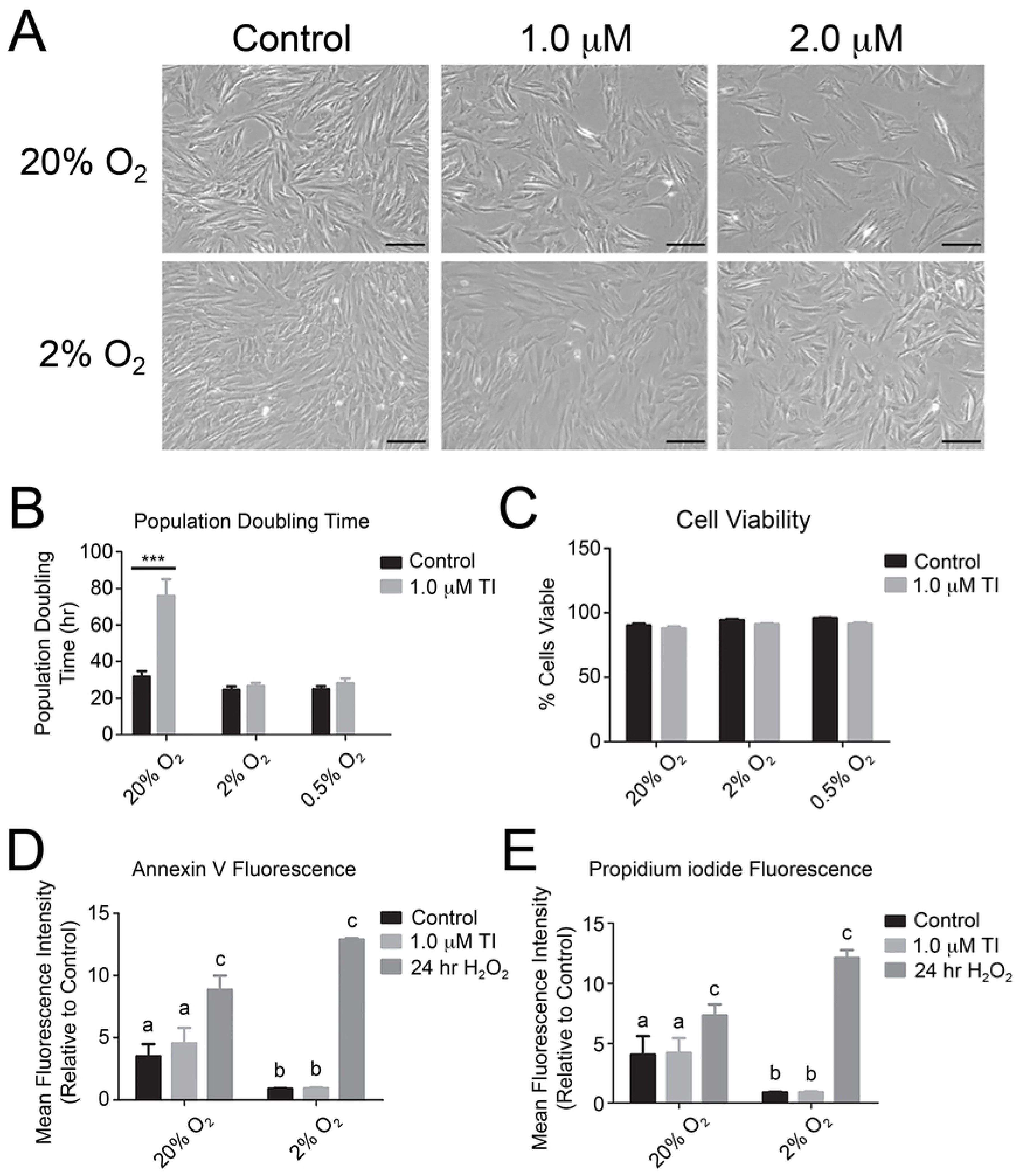
Telomerase inhibition causes cell growth arrest of fetal guinea pig muscle cells without indication of apoptosis. (A) Fetal guinea pig muscle cells grown under 20% O_2_ conditions and treated with telomerase inhibitor (TI-IX) exhibited reduced confluency and altered cell morphology typical of cells in senescence. Scale bars represent 100 μM. (B) Cells were seeded at different concentrations and counted every 24 hours for 3 days to determine population-doubling times in different oxygen concentrations and treatment with 1 μM TI-IX. There was a significant increase in population doubling time for 20% O_2_ cultured cells treated with TI-IX (***; p<0.001). (C) Cells were cultured for 48 hr with or without TI-IX in 20% O_2_, 2% O_2_ and 0.5% O_2_ and then stained with trypan blue exclusion to determine cell viability. All results were analyzed with a two-way ANOVA, n=3. (D and E) Cells were incubated for 48 hr at 20% and 2% O_2_ with or without TI-IX with a positive control of a 24 hr treatment with 500 µM H_2_O_2_. Cells were trypsinized and stained for extracellular apoptotic marker Annexin V, and an intracellular fluorescent marker propidium iodide (PI) to label apoptotic/necrotic cells. The cells were analyzed in the Accuri C6 flow cytometer, and mean fluorescence of each stain was determined. Mean fluorescence was graphed and shown for Annexin V (D), and for PI (E). Results were analyzed with two-way ANOVA, n=4. Labelled bars with the same letter were not significantly different (p>0.05) from each other. Labelled bars with a and b indicate a significant difference of p<0.05, and c labelled bars indicate a significant difference of p<0.01 compared to a and b.

To strengthen the observed senescent phenotype, cells were seeded into 6-well dishes at different concentrations of 25,000, 35,000 and 50,000 cells and were counted every 24 hours for 3 days to determine population-doubling time. There was no significant (p>0.05) difference between cells grown at 2% O_2_ and 0.5% O_2_ with TI-IX treatment, but there was a significant increase (p<0.001) in population doubling time for fetal guinea pig muscle cells grown at 20% O_2_ and treated with 1 μM TI-IX (Fig 2B). Cell viability was determined after 48 hours in both control and telomerase inhibited cells using trypan blue and hemocytometer counts. There was no significant difference (p>0.05) in fetal muscle cell viability at different oxygen tensions or Telomerase Inhibitor IX concentrations (Fig 2C).

Annexin V and propidium iodide (PI) staining were used to determine if telomerase inhibition triggered apoptosis in fetal guinea pig muscle cells. There was no significant difference between cells cultured at the same oxygen concentration, however cells cultured at 20% O_2_ tensions had significantly (p<0.05) higher Annexin V fluorescence and PI staining compared to cells cultured in 2% O_2_ conditions (Fig 2D and E). Annexin V Fluorescence and PI staining of the positive control cells (H_2_O_2_ treated) were significantly (p<0.01) higher compared to non-treated control and Telomerase Inhibitor treated cells (Fig 2D and E).

### Telomerase inhibition causes alteration in tumor suppressor proteins that signify cell cycle arrest

DNA damage-induced activation of ATM/ATR and Chk1/Chk2 phosphorylates the tumor suppressor protein p53, which affects transcription of many downstream proteins including the cell cycle inhibitor p21. Together p53 and p21 mediates cellular senescence when cells are exposed to certain stressors and damage [67]. With the addition of Telomerase Inhibitor IX, p53 protein levels were significantly elevated in fetal muscle cells cultured in 20% O_2_ (p<0.05), 2% O_2_ (p<0.01) and 0.5% O_2_ (p<0.05) atmospheres (Fig 3A). Increases in p53 protein were accompanied with elevated phospho-p53 at serine-15, but these increases were not significant (Fig 3B). Following the expression levels of p53, p21 was significantly increased two-fold at each oxygen concentration with Telomerase Inhibitor IX treatment of muscle cells cultured in 20% O_2_ (p<0.001), 2% O_2_ and 0.5% O_2_ (p<0.05) conditions (Fig 3C). Retinoblastoma protein (Rb) is a critical regulator of the G1/S checkpoint and must be phosphorylated (pRb) in order to be inactivated and release of the E2F transcription factors to allow the cell to continue through the cell cycle. There was no difference between any of the groups when immunoblotting for non-phosphorylated Rb protein levels (Fig 3E). However, cells cultured in 20% O_2_ and treated with Telomerase Inhibitor IX showed significantly decreased (p<0.05) levels of phosphorylated (S795) Rb protein (Fig 3F), which decreased more significantly (p<0.001) when the ratio of pRb to total Rb levels were calculated (Fig 3G). Cells cultured in 2% O_2_ displayed no significant difference (p>0.05) between non-treated controls and inhibitor treated cells.

**Fig 3.**
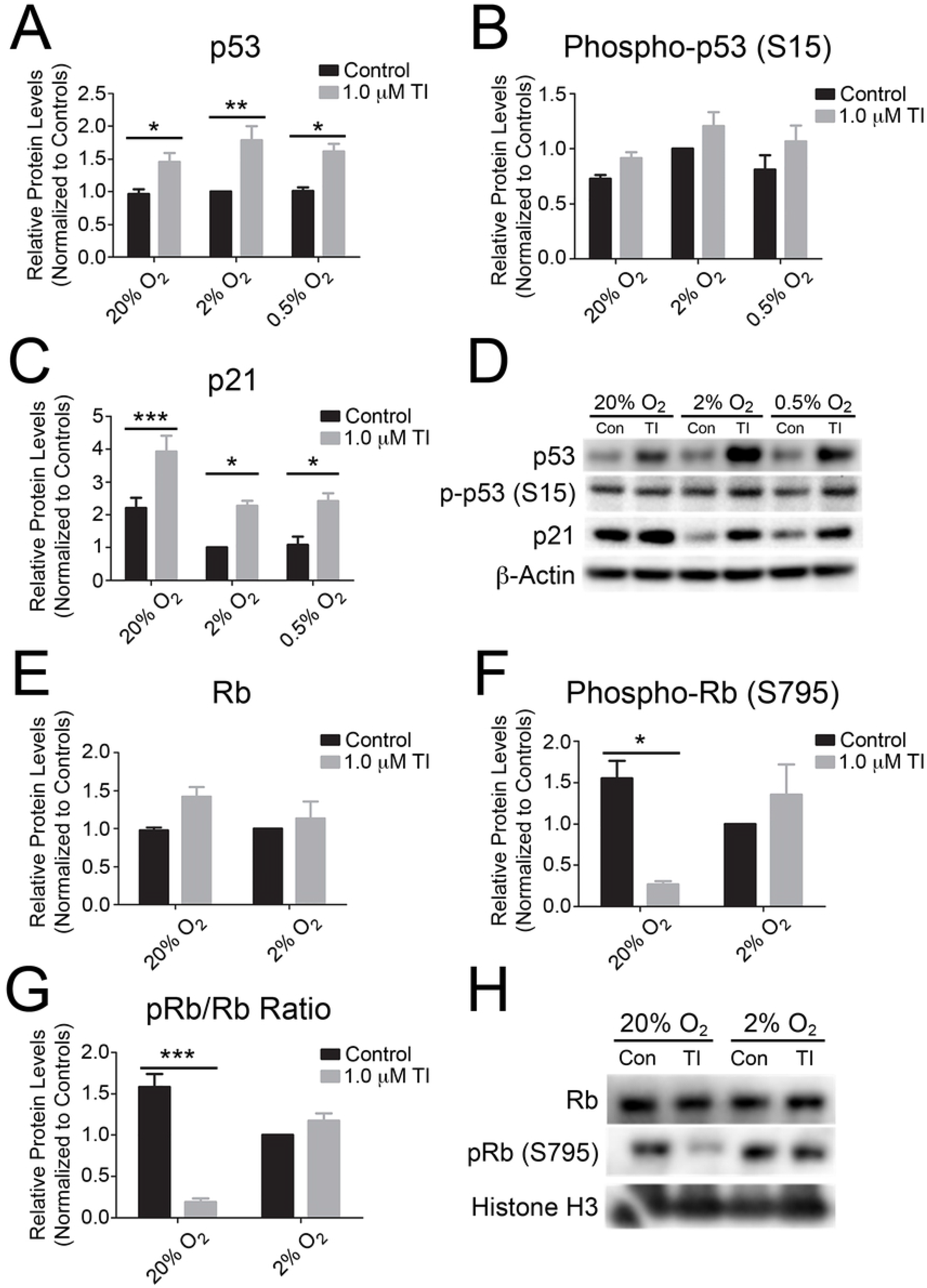
Elevated senescence marker expression in fetal guinea pig muscle cells treated with telomerase inhibition. Fetal guinea pig muscle cells were treated for 48 hr at 20%, 2%, and 0.5% oxygen (O_2_) tensions with or without Telomerase Inhibitor (TI-IX). Total protein extractions were immunoblotted followed by densitometric analyses of blots exposed to antibodies specific to (A) p53, (B) phospho-p53 (Ser15), and (C) p21. β-actin protein levels were detected as a loading control. A representative western blot is shown in (D). Results were analyzed with a two-way ANOVA, n=5, with (***), (**) and (*) indicating p<0.001, p<0.01, and p<0.05, respectively. (E-H) Cells were collected and nuclear protein extractions were used in immunoblotting followed by densitometric analyses of blots exposed to antibodies specific to (E) Rb, and (F) phospho-Rb (Ser795). A ratio of pRb/Rb was calculated to determine amount of total Rb that was phosphorylated (G). Total histone H3 protein levels were detected as a loading control. Results were analyzed with a two-way ANOVA, n=3, with (***) and (*) indicating p<0.001, and p<0.05, respectively. A representative Western blot is shown in (H).

### Significant increase in telomere-damage induced foci in telomerase inhibited fetal guinea pig muscle cells

Telomerase has been shown to play a role in the DNA damage response, and its inhibition may be a potential reason for the increase in cell senescence markers. To determine if there is an increased amount of DNA damage, the DNA double strand break marker phosphorylated ɣ-H2AX was assayed by immunoblotting and immuno-FISH with a telomere DNA probe. Total protein levels of phospho-ɣH2AX showed a slight increase when treated with Telomerase Inhibitor IX, and were increased as the oxygen concentration decreased (Fig 4C). However, densitometric analysis of the immunoblots revealed that these differences were not significant (p>0.05). Telomere induced foci (TIFs) are comprised of co-localization of DNA damage markers, such as phospho-ɣH2AX with a fluorescent telomeric probe (Fig 4A). A significant increase in TIF-positive cells was observed in TI-IX treated cells at both 20% O_2_ (p<0.01) and 2% O_2_ (p<0.05) tensions compared to non-treated controls (Fig 4B). Furthermore, there was greater TIF-positive cells for cells grown under 20% O_2_ conditions compared to those cultured in 2% O_2_ (p<0.05).

**Fig 4.**
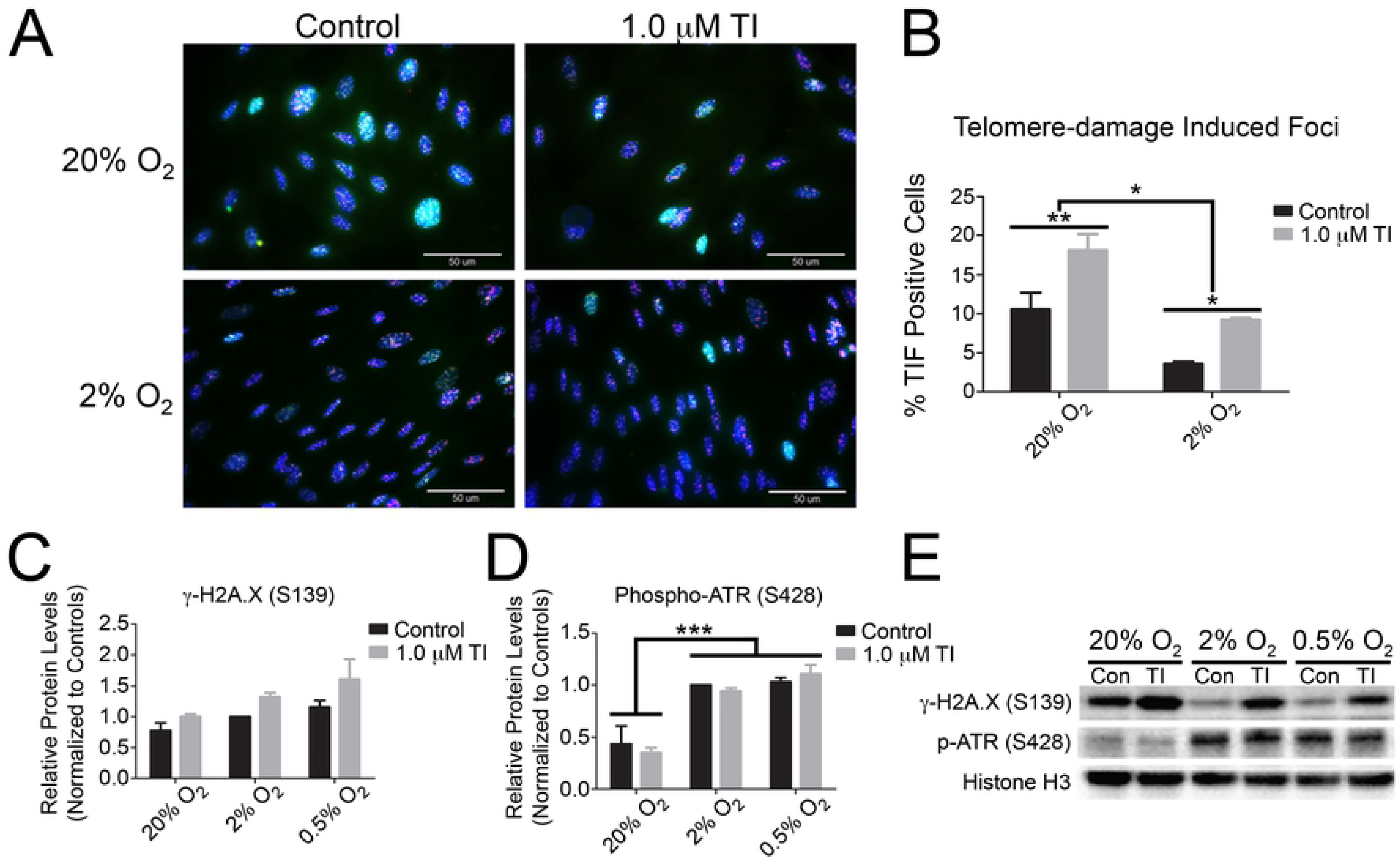
Increased frequency of telomere-damage induced foci in telomerase inhibited fetal guinea pig muscle cells. Fetal guinea pig muscle cells were cultured in chamber slides at 20% and 2% oxygen (O_2_) with or without Telomerase Inhibitor (TI-IX). Cells were fixed and labelled with a primary antibody for phospho-ɣH2AX, and secondary antibody conjugated with Alexa-Fluor 488 to identify sites of DNA damage. Fluorescent *in situ* hybridization of a Cy5 fluorescent tagged Telomere PNA probe allowed signals to be localized to the telomere as telomere dysfunction induced foci (TIF). Co-localizations of these signals with Hoescht 33342 nuclear staining are shown in (A), with scale bars representing 50 μM. The number of TIF-positive cells were counted and represented in (B). Significant differences between groups were calculated with a two-way ANOVA, n=4, with (*) and (**) representing p<0.05 and p<0.01, respectively. (C-E) Altered DNA damage response proteins ɣH2AX and ATR. Fetal guinea pig muscle cells were treated for 48 hours at 20%, 2%, and 0.5% O_2_ with or without TI-IX. Cells were collected and total protein extractions were used in immunoblotting followed by densitometric analyses of blots exposed to antibodies specific to (C) phospho-ɣH2AX (Ser139), and (D) phospho-ATR (Ser428). Total histone H3 protein levels were detected as a loading control. A representative western blot is shown in (E). Results were analyzed with a two-way ANOVA, n=3, with (***) indicating p<0.001.

The DNA damage response is initially sensed at the DNA level by the MRN complex and the ATM/ATR proteins. ATM and ATR are kinases that phosphorylate many downstream targets involved in cell cycle arrest during DNA damage and DNA repair proteins. To determine if ATR was active, a phospho-ATR (S428) antibody was utilized for immunoblotting. There were no significant differences between control and cells treated with Telomerase Inhibitor IX, but there was a significant increase in phospho-ATR (S428) levels within fetal muscle cells cultured at lower oxygen concentrations compared to those at 20% O_2_ (p<0.001) (Fig 4D).

### Disturbed intracellular redox homeostasis in high oxygen cultured fetal guinea pig muscle cells treated with telomerase inhibition

Due to the increased telomeric DNA damage observed in cultured fetal muscle cells treated briefly with telomerase inhibitor, mitochondrial ROS was evaluated due to its known involvement in regulating the senescence state [29]. MitoTracker® Red CMXRos is a red-fluorescent dye that fluoresces upon oxidation in live cells and is sequestered into mitochondria based on its membrane potential. Guinea pig muscle cells displayed red fluorescence in the mitochondria, shown as small punctate points surrounding the nucleus (Fig 5A). Flow cytometric analysis of MitoTracker fluorescence was significantly (p<0.001) increased in cells cultured in 20% O_2_ compared to those grown in 2% O_2_ (Fig 5B). MitoTracker® Red CMXRos levels were significantly increased by telomerase inhibition at 20% O_2_ (p<0.05) but not at 2% O_2_ conditions (Fig 5B).

**Fig 5.**
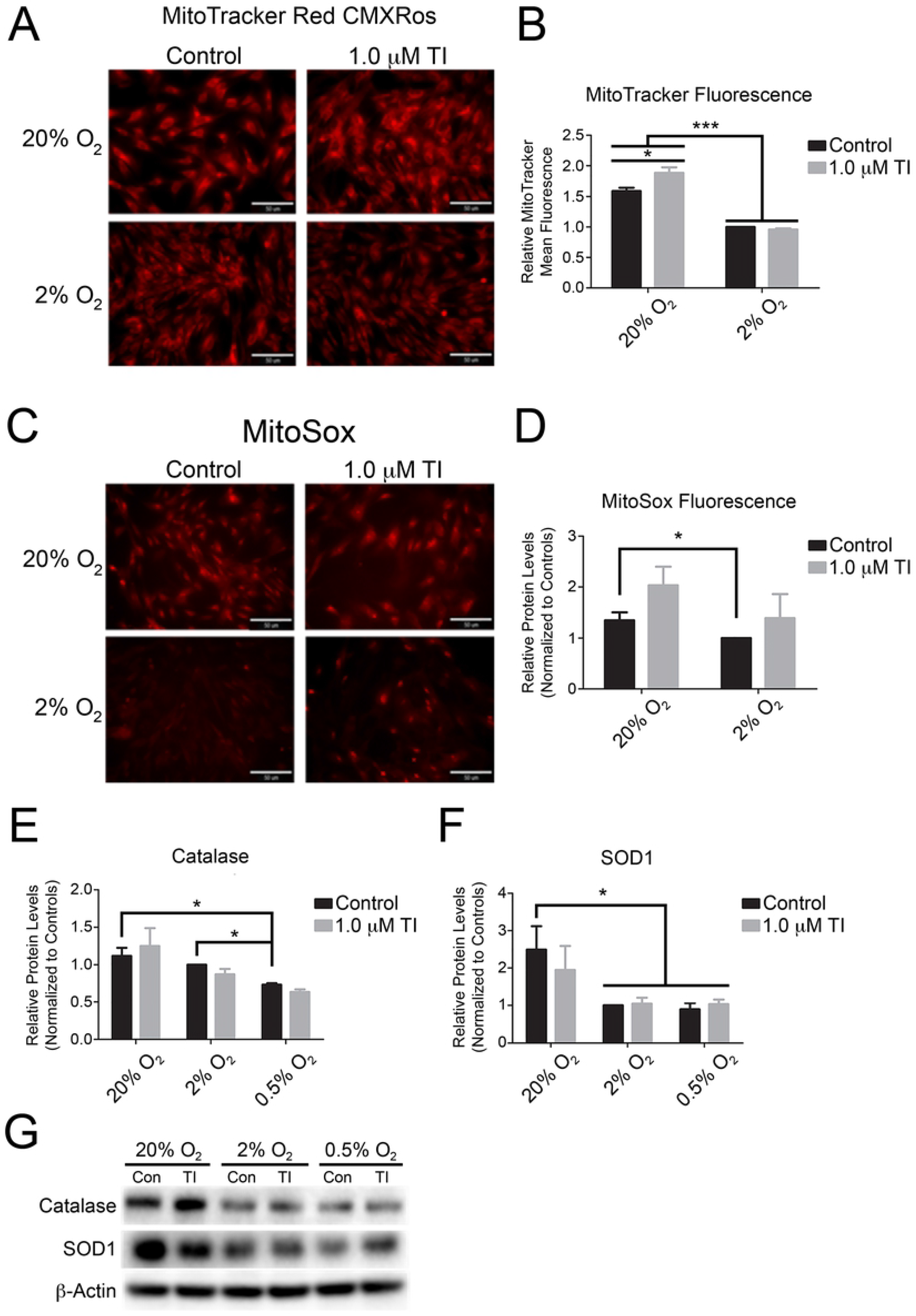
Elevated mitochondrial membrane potential and superoxide production despite higher antioxidant levels in telomerase inhibited fetal guinea pig muscle cells cultured at 20% oxygen tensions. MitoTracker® Red CMXRos or MitoSOX™ Red was incubated with fetal guinea pig muscle cells for 15 min following a 48 hr treatment with or without Telomerase Inhibitor (TI-IX) at 20% and 2% O_2_ conditions. Cells were then visualized by fluorescent microscopy (A and C) or analyzed with the Accuri C6 flow cytometer with mean fluorescence determined and represented graphically in (B and D). Results were analyzed with a two-way ANOVA, with n=3 for MitoTracker® Red CMXRos and n=4 for MitoSOX™ Red experiments. Significant differences are indicated by (*) and (***), representing p<0.05, and p<0.001, respectively. Scale bars represent 50 μM. (E-G) Antioxidant proteins catalase and SOD1 are elevated in fetal muscle cells cultured in higher oxygen concentrations. Fetal muscle cells were treated for 48 hours at 20%, 2%, and 0.5% O_2_ with or without TI-IX. Total protein extractions were used in immunoblotting followed by densitometric analyses of blots exposed to antibodies specific to catalase (E) and SOD1 (F). Total β-actin protein levels were detected as a loading control. A representative western blot is shown in (G). Results were analyzed with a two-way ANOVA, n=3, with (*) indicating p<0.05.

MitoSOX is used as a mitochondrial superoxide indicator, which when oxidized will fluoresce red when bound to nucleic acids. Guinea pig muscle cells were incubated with MitoSOX and then used in flow cytometry to analyze mean MitoSOX fluorescence (Fig 5C). Red fluorescence was mainly localized to the nucleus, but at higher oxygen concentrations showed some diffuse fluorescence surrounding the nucleus in the cytoplasm. Flow cytometric analysis revealed trends indicating an increase in fluorescence with telomerase inhibition but these were not significant (p>0.05), however cells grown in 20% O_2_ did display significantly (p<0.05) elevated superoxide levels compared to those cultured in 2% O_2_ conditions (Fig 5D).

The general oxidative stress indicator 6-carboxy-2’,7’-dichlorodihydrofluorescein diacetate (carboxy-H_2_DCFDA) was used to assess cytoplasmic ROS (primarily H_2_O_2_). Fetal muscle cells were incubated with carboxy-H_2_DCFDA and mean fluorescence intensities analyzed by flow cytometry (S2 Fig). Like other oxidative stress indicators, carboxy-H_2_DCFDA fluorescence was significantly higher in 20% O_2_ compared to 2% O_2_ (p<0.01). A unique finding was that telomerase inhibition was able to induce a significant decrease in carboxy-H_2_DCFDA fluorescence in 20% O_2_ cultured cells (p<0.05) and a non-significant (p>0.05) reduction in 2% O_2_ cultured cells. Carboxy-H_2_DCFDA fluorescence was shown to have diffuse green fluorescence throughout the cell (not shown).

The antioxidant proteins SOD1 and catalase were also evaluated by immunoblotting. SOD1 is a soluble cytoplasmic and mitochondrial intermembrane space enzyme responsible for converting superoxide radicals into molecular oxygen and hydrogen peroxide (H_2_O_2_), while catalase is the major enzyme for converting H_2_O_2_ into water and oxygen in a cell. Catalase levels were significantly higher (p<0.05) in both 20% O_2_ and 2% O_2_ cultured cells than cells grown in 0.5% O_2_ (Fig 5E, G). Catalase levels were also at their highest in fetal muscle cells grown in 20% O_2_ and was decreased in lower oxygen concentrations. Levels of SOD1 protein was significantly increased (p<0.05) in 20% O_2_ cultures compared to cells grown in 2% O_2_ and 0.5% O_2_ (Fig 5F, G). There was no significant difference in the levels of both catalase and SOD1 following telomerase inhibition compared to non-treated cells (Fig 5E-G).

### Altered metabolic signaling in telomerase inhibited fetal guinea pig muscle cells

The mTOR pathway is necessary for integrating different stimuli and regulating cell growth and metabolism, and may play a role in triggering senescence [68]. Immunoblotting for mTOR and phopsho-mTOR revealed no difference in mTOR protein levels between treatment groups, nevertheless there was a significant decrease in phospho-mTOR at 20% O_2_ (p<0.05) compared to cells grown in both 2% O_2_ and 0.5% O_2_ conditions (Fig 6A, B). However, phosphorylated mTOR and mTOR protein ratios did not reveal any significant differences (p>0.05) among groups (Fig 6C).

**Fig 6.**
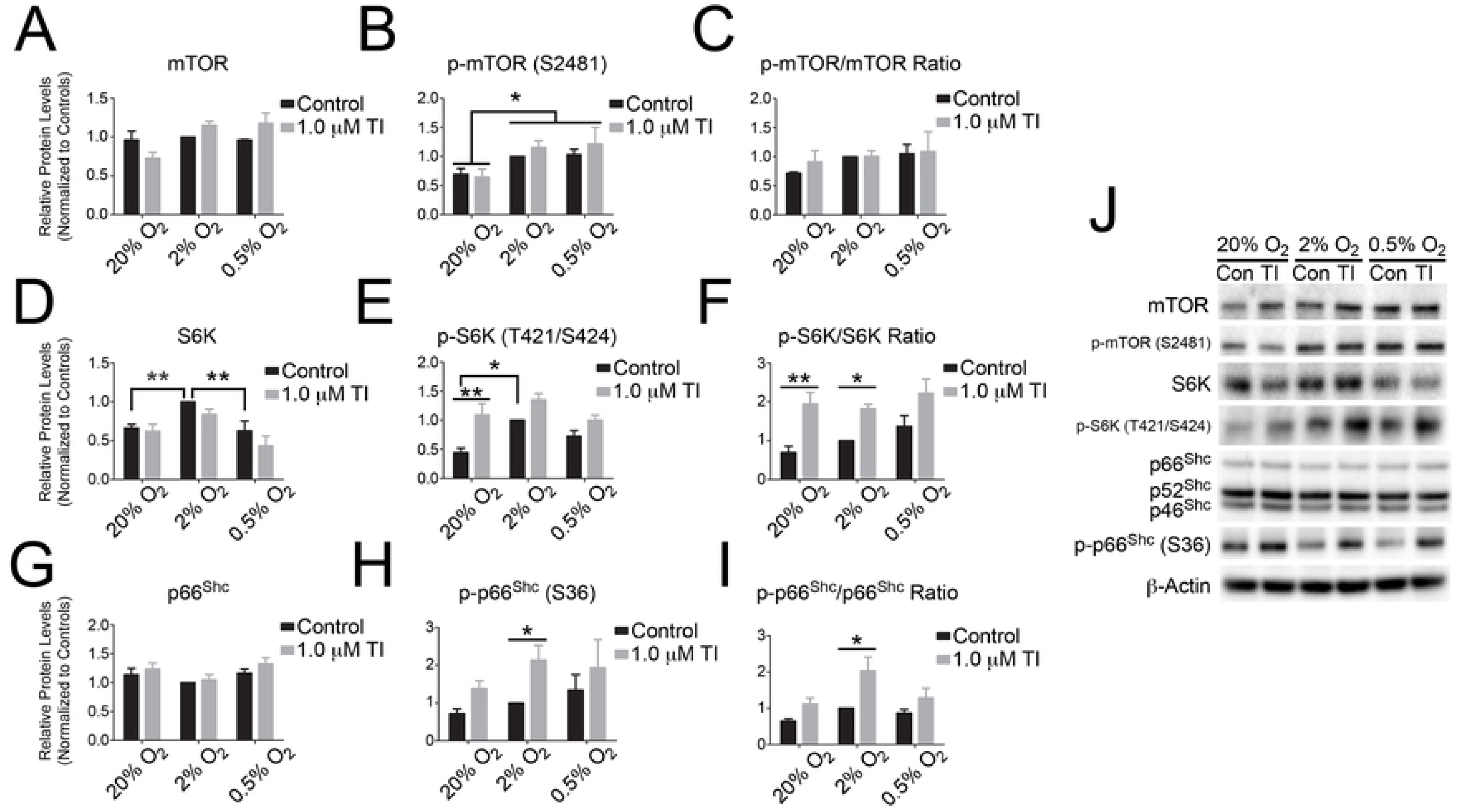
The stress adaptor protein p66Shc is associated with S6K activation in telomerase inhibited fetal guinea pig muscle cells. Fetal guinea pig muscle cells were treated for 48 hours at 20%, 2%, and 0.5% oxygen tensions with or without 1 μM Telomerase Inhibitor IX. Cells were collected and total protein extractions were used in immunoblotting with antibodies specific to mTOR, p-mTOR (S2481), S6K, p-S6K (Thr421/Ser424), p66Shc and p-p66Shc (S36) and β-actin as a loading control. Densitometry was performed and represented for mTOR (A), p-mTOR (B), S6K (D), p-S6K (E), p66Shc (G) and p-p66Shc (H). The densitometries of phosphorylated to total protein ratios were calculated and shown (C, F, I). A representative western blot is shown in (J). Results were analyzed with a two-way ANOVA, n=4, with (**) and (*) indicating p<0.01, and p<0.05, respectively.

S6K is the downstream kinase in the mTOR pathway that regulates ribosomal protein synthesis. Total S6K protein levels were increased in cells grown at 2% O_2_ compared to cells grown in 20% O_2_ and 0.5% O_2_ (p<0.01), but was not different with telomerase inhibition (Fig 6D). Phosphorylated S6K (T421/S424) was not significantly (p>0.05) up regulated between control cells and cells treated with the telomerase inhibitor at both 2% and 0.5% O_2_, but was significantly (p<0.01) elevated in 20% O_2_ cultures (Fig 6E). Similar to S6K, phospho-S6K was significantly (p<0.05) decreased in 20% O_2_ control cells compared to fetal muscle cells cultured in 2% O_2_ (Fig 6E). There were significant increases in phospho-S6K/S6K ratios at both 20% O_2_ and 2% O_2_ (p<0.01) tensions after telomerase inhibition, and was up regulated at 0.5% O_2_ although this increase was not significant (p>0.05; Fig 6F).

P66Shc is a unique adaptor protein as it has been reported to have roles in both sensing oxidative stress, mediating its effects and even causing more ROS to be generated at the mitochondrial level [69]. Thus, due to this role in ROS sensing/production, it has been implicated as an integral regulator of cellular aging by inducing apoptosis and senescence [69, 70]. Interestingly, there was an overall trend for increasing phosphorylated (S36)-p66Shc levels in fetal muscle cells grown at lower oxygen concentrations. Phosphorylated (S36) p66Shc and the p-p66Shc/p66Shc ratios were increased at all oxygen concentrations after treatment with Telomerase Inhibitor IX, but this was only significant (p<0.05) at 2% O_2_ conditions (Fig 6H, I).

### Akt/GSK3 signaling is altered in telomerase-inhibited cells

Akt is a major regulator on cell growth and metabolism, and can phosphorylate mTOR and activate downstream pathways involved in cell proliferation [71]. Akt protein levels, determined using a pan Akt antibody, were significantly decreased in fetal muscle cells cultured in 20% O_2_ with telomerase inhibition (p<0.05) compared to untreated control cells (Fig 7A, F). Immunoblotting with a phospho-Akt (T303) antibody exhibited the same significant decrease at 20% O_2_ and with telomerase inhibition (p<0.05; Fig 7B, F). However, the ratio of phosho-Akt/Akt protein levels did not show any significant differences between any groups (p>0.05; Fig 7C, F).

**Fig 7.**
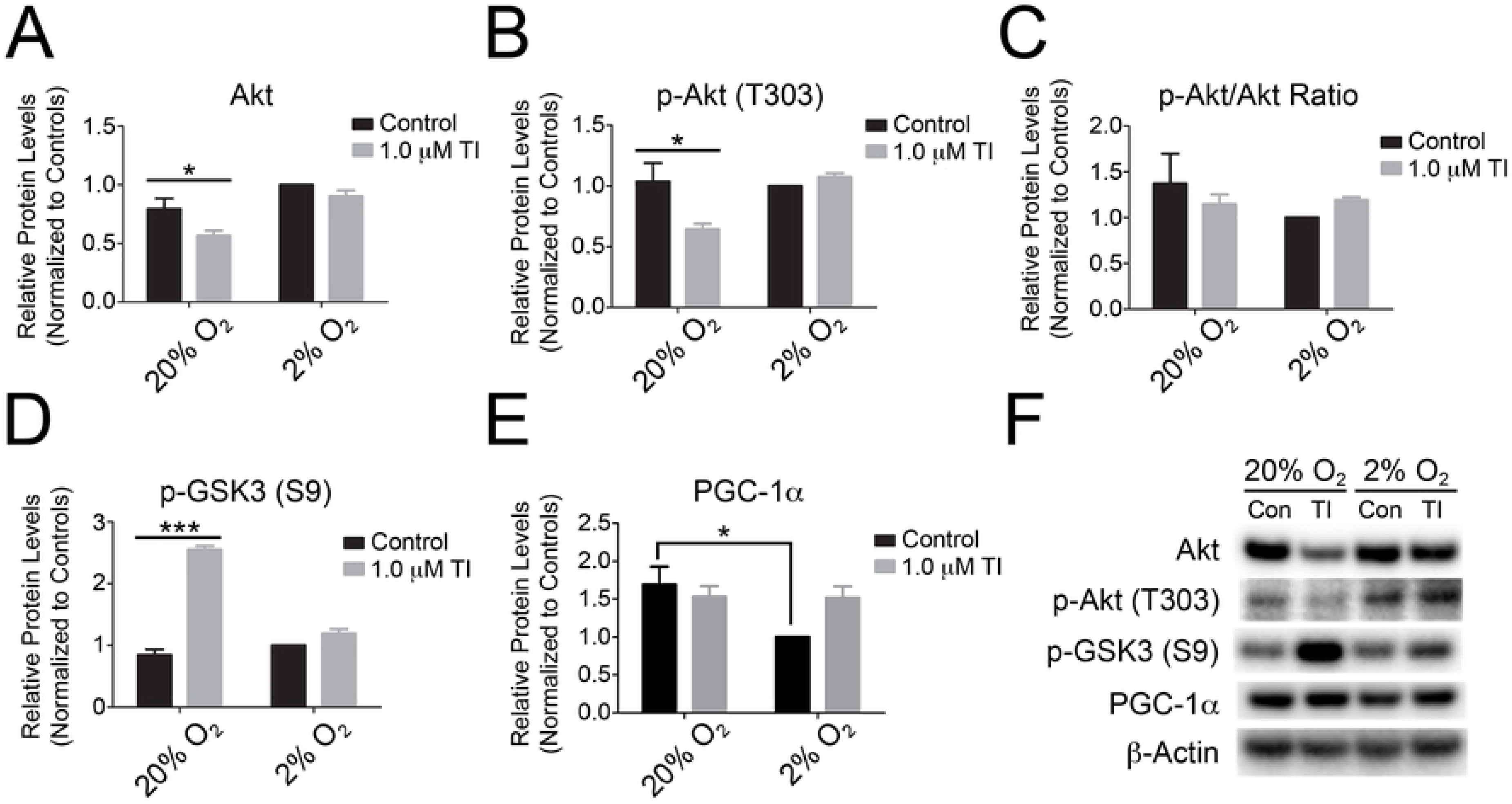
Altered Akt/GSK3 signaling in telomerase-inhibited fetal guinea pig muscle cells cultured under high oxygen. Fetal guinea pig muscle cells were treated for 48 hours at 20%, and 2% oxygen with or without Telomerase Inhibitor IX. Cells were collected and total protein extractions were used in immunoblotting with antibodies specific to pan Akt, phospho-Akt (Thr308), p-GSK3 (S9), PGC-1α and β-actin as a loading control. Densitometry was performed and represented for pan Akt (A), phospho-Akt (Thr308) (B), p-GSK3 (D), PGC-1α (E). The densitometries of phosphorylated-Akt to total Akt protein ratio was calculated and shown (C). A representative western blot is shown in (F). Results were analyzed with a two-way ANOVA, n=3, (*) indicating p<0.05 and (***) indicating p<0.001.

GSK-3 is involved in the PI3K/Akt pathway and is therefore also involved in cell proliferation and may be involved in the regulation of senescence and apoptosis [72]. GSK-3 is inactive when phosphorylated at serine 9 and serine 21. A significant (p<0.001) increase in phospho-GSK-3 at serine 9 was observed in GPMCs treated with TI-IX at 20% O_2_ (Fig 7D, F). There were no differences in phospho-GSK-3 levels between cells incubated at 2% O_2_ with and without telomerase inhibition.

PGC-1α is a transcriptional co-activator that regulates many genes in energy metabolism and mitochondrial biogenesis [73]. PGC-1α was recently linked to telomere dysfunction through p53 activation [49]. However, no difference in PGC-1α levels were observed between control and telomerase inhibited cells. PGC-1α was significantly (p<0.05) higher in 20% O_2_ untreated cultured cells compared to their 2% O_2_ untreated counterparts, which may indicate an increase mitochondrial biogenesis in cells cultured at higher oxygen concentrations (Fig 7E, F).

## Discussion

This study examined the effects of telomerase inhibition on signaling pathways related to cellular senescence, DNA damage and metabolism in fetal guinea pig muscle cells. Cells cultured under various oxygen tensions and treated with Telomerase Inhibitor (TI)-IX displayed significantly decreased telomerase activity levels. Significant telomerase inhibition was observed in cells grown in high (20%) oxygen concentrations compared to those cultured under low (2% and 0.5%) oxygen conditions. Telomerase inhibition induced senescence as demonstrated by increased levels of senescent markers p53 and p21, decreased phosphorylation of the cell cycle regulator Rb protein and prolonged cell cycle of cells cultured under 20% oxygen conditions. Senescence detection was correlated to an increase in DNA damage along with a significant increase in the number of cells displaying telomere dysfunction induced foci (TIFs), which are an indication of telomere uncapping. Telomerase inhibition increased mitochondrial ROS production, however, overall intracellular ROS levels were decreased. Activated p66Shc levels were increased upon telomerase inhibition, suggesting a mechanism for the higher levels of mitochondrial ROS and DNA damage. Furthermore, proteins involved in nutrient sensing, cell proliferation and metabolism pathways showed a down regulation with telomerase inhibition including Akt, phospho-Akt and GSK-3. Phosphorylated mTOR levels were not increased by telomerase inhibition, however its downstream effector kinase S6K showed a significant increase in phosphorylation/activation. Interestingly, one of the key master regulators of cellular metabolism, peroxisome proliferator-activated receptor gamma co-activator 1-alpha (PGC-1α) was increased in cells grown under high oxygen concentrations.

The specific mechanism of action of TI-IX is currently unclear, but it may inhibit telomerase by transcriptional and translational inhibition of telomerase reverse transcriptase (TERT), like its parent molecule epigallocatechin-3-gallate (EGCG) [74, 75]. Previous reports using TI-IX and other small molecule telomerase inhibitors led to telomere shortening and senescence of various normal somatic and cancer cell lines [76–79]. RNAi-mediated knockdown of TERT in hepatocellular carcinoma cells [80] and breast cancer cells [81] increased their sensitivity to apoptosis. Interestingly, in our study non-treated fetal muscle cells incubated in 2% O_2_ displayed significantly higher telomerase activity compared to cells cultured in 20% O_2_. Previous reports have shown that low oxygen tensions can upregulate telomerase activity [82–84] and that hypoxia inducible factor (HIF-1α) can increase transcription of TERT mRNA [83, 85]. Due to the upregulation of telomerase in low oxygen cultures, treatment of cells with TI-IX at 2% O_2_ was not able to decrease telomerase activity by the same level as cells treated in 20% O_2_ conditions. A higher dose of TI-IX was required to significantly reduce telomerase activity levels in 2% O_2_ cultured cells compared to cells grown in 20% O_2_. Low oxygen cells not only contained higher telomerase activity, they appeared to be more resistant to the effects of telomerase inhibition, suggesting that fetal muscle cells cultured under these conditions have a greater ability to maintain telomerase activity levels and be resistant to senescence and apoptosis.

TI-IX treatment altered cellular morphology, and this was more apparent at 20% O_2_ compared to cells treated under 2% O_2_ conditions. Cells appeared larger, flatter and were less confluent after telomerase inhibition. The observed morphological changes were consistent with those of other senescent cell types [76, 86]. The population doubling time for TI-IX treated cells was found to be significantly longer in 20% O_2_ cultured cells supporting the decreased cellular confluency and growth arrest after telomerase inhibition. Unlike cells grown at 20% O_2_, there were only marginal differences in population doubling time observed in cells cultured at lower oxygen concentrations with telomerase inhibition, and no significant differences in population doubling time were measured in control cells between oxygen concentrations, further supporting that 20% oxygen conditions are suboptimal.

Since telomerase inhibition may also trigger apoptosis [80, 81], markers of apoptosis were analyzed by flow cytometry to strengthen the cellular growth arrest phenotype. The incidence of apoptosis did not increase upon telomerase inhibition since both Annexin V and PI fluorescence levels did not differ significantly between control cells and those treated with TI-IX. However, there was a significant increase in apoptotic cells incubated at 20% O_2_ compared to those at 2% O_2_ further supporting that atmospheric oxygen is inherently a more stressful *in vitro* environment that can lead to cellular apoptosis/senescence when cells are exposed to another challenge such as telomerase inhibition. Importantly, cell viability between oxygen concentrations and treatments was not significantly affected. Since telomerase inhibited fetal muscle cells cultured in 20% oxygen conditions displayed morphological characteristics of senescence without an increase in apoptosis, we next examined the levels of various molecular markers of cellular senescence.

Telomerase may have a role in DNA damage and repair, and disruption of these pathways can lead to cellular senescence [87, 88]. At all oxygen concentrations, we observed a significant increase in p53 accumulation with telomerase inhibition. p53 accumulation is regulated by its phosphorylation, and when phosphorylated at serine-15 and serine-20 can bypass its targeting for degradation [89]. The measured increase in p53 protein levels following telomerase inhibition indicates that this is likely occurring, even though the phospho-p53 (Ser15) immunoblotting was not significantly increased. Other phosphorylation sites (e.g. serine-20) may have been significantly up regulated but these were not investigated. One of the main targets of p53 in response to DNA damage is the cell cycle inhibitor, p21 [90]. We measured a significant two-fold increase in p21 upon telomerase inhibition at all oxygen concentrations examined, although it was initially higher in non-treated cells cultured at 20% O_2_ compared to those at 2% O_2_ and 0.5% O_2_ tensions. p21 halts the cell cycle through its inhibition of cyclin-dependent kinases and their counterpart cyclins, which decreases phosphorylation of Rb [91]. Immunoblotting for phosphorylated Rb verified that hypophosphorylation only occurred in 20% O_2_ telomerase inhibited cells, which associated strongly with the longer population doubling time measured for this cell group. Previously, Dyskeratosis Congenita cell lines have shown increased p53 and p21 levels in response to telomerase deficiency and expression levels of each positively correlated with the oxygen concentration of the cell cultures [92]. P53 was found to be necessary for the progression of senescence in an irradiated breast tumor cell line, and without functional p53 these cells could only activate apoptosis [93]. The expression of transfected p21 alone in glioma cells triggers permanent cell cycle arrest and may itself have a role in decreasing telomerase activity levels [94]. Oxygen concentration may also play a role, as 20% O_2_ cultured cells had higher levels of p21, with and without telomerase inhibition, as induction of p21 could be due to hyperoxic oxygen conditions through stimulation of p53. Conversely, telomerase overexpression in human embryonic stem cells have shown hyperphosphorylation of Rb and decreased G1 phase of the cell cycle [95]. These studies in conjunction with our results reveal that senescence can be initiated through decreases in telomerase activity and maintained through activation of the p53-p21-Rb pathway.

DNA damage including telomeric DNA damage can induce an increase in senescent signaling pathway markers [65, 88]. To further investigate the role of telomerase inhibition in DNA damage, the marker phospho-γ-H2AX was used to quantify the amount of DNA double strand breaks throughout the genome and at telomeres. There was an increase in total phospho-γ-H2AX protein levels with telomerase inhibition at all oxygen concentrations. This DNA damage was shown to be localized to the telomeres by co-localization of phospho-γ-H2AX at the telomeres by immuno-FISH. The number of telomere-damage induced foci (TIF)-positive cells were elevated with telomerase inhibition at both 20% O_2_ and 2% O_2_, and that cells at 20% O_2_ also had significantly more TIF-positive cells compared to those cultured at 2% O_2_. Telomere uncapping, recognized by TIFs [96], are sites of DNA damage that induce senescence [97, 98]. Telomeres are favored targets for DNA damage in aging and stressed tissues [99], and increases in telomeric DNA damage have been correlated with decreased telomerase activity [100]. Under severe hyperoxic conditions (40% O_2_), DNA damage and telomere shortening is increased due to active telomerase translocating from the nucleus into the mitochondria [39]. These studies are in agreement with our own findings showing increased telomeric DNA damage in cells with decreased telomerase activity under higher levels of oxygenation.

The DNA damage response protein ATR was investigated to determine if the DNA damage response pathway was activated. The addition of TI-IX showed no significant differences for phosphorylated ATR, however levels were significantly lower in 20% O_2_ fetal muscle cells than those cultured under lower oxygen concentrations. Phospho-ATR has been reported to be up regulated in hypoxic conditions in tumors, and our result supports that ATR activity may be oxygen-dependent [101]. This increase in phospho-ATR may be related to proliferation rates, and may aid cells cultured at lower oxygen levels in coping with telomerase inhibition by repairing DNA damage instead of becoming senescent [102]. ATR may not play an important role in the response to DNA damage in our model since it was not activated with higher levels of DNA damage at 20% O_2_, but instead could display higher activity through activation of DNA replication damage in cells with a faster population doubling time [102, 103]. If there is excessive DNA damage, it may be irreparable, which may mitigate the DNA damage response [104]. We did not observe increased phospho-ATR with telomerase inhibition and DNA damage, but this could be due to activations of other DNA damage response proteins, such as ATM [105].

Telomerase is known to shuttle between the nucleus, cytoplasm and mitochondria in cells, but its exact function in mitochondrial and cytoplasmic compartments are not well recognized [106]. In stress conditions, up to 80-90% of telomerase can shuttle out of the nucleus into the mitochondria, resulting in telomere shortening/damage [39]. Telomerase may have a role in mitochondrial DNA protection or to alleviate mitochondrial ROS production [39, 40]. Similar to previous studies [39, 65, 106], we measured an increase in mitochondrial ROS in telomerase inhibited 20% O_2_ cells. This ROS increase was not observed in 2% O_2_ cultured cells with telomerase inhibition. This suggests a role for enzymatically active telomerase in preventing mitochondrial ROS production, increasing mitochondrial membrane potential or improving mitochondrial coupling [39, 40, 106]. The increased mitochondrial ROS observed in telomerase inhibited 20% O_2_ cells may also be due to an increase in p53, which modulates expression of redox-related genes involved in mitochondrial permeability [107–109]. We observed an increase in cytoplasmic/total cellular ROS in cells cultured under 20% O_2_ compared to those cultured in 2% O_2_ further supporting a hyperoxic environment for cells grown at atmospheric O_2_ tensions. Surprisingly, we detected decreased cytoplasmic ROS levels in telomerase inhibited cells. Contrary to reports that telomerase improves mitochondrial function [39], mitochondrial TERT also appears to exacerbate free-radical-mediated mtDNA damage [110]. These results support differing roles for telomerase in regulating ROS production, but further research is needed to fully elucidate its role in the mitochondria.

The transcription factor PGC-1α regulates genes involved in energy metabolism and mitochondrial biogenesis [73], and was significantly increased in 20% O_2_ compared to 2% O_2_ cells. Elevated PGC-1α correlated with increased mitochondrial number in cells grown at 20% O_2_ and could be responsible for increased ROS levels [111]. Coupled with results indicating higher levels of DNA damage and senescent markers, 20% O_2_ promotes a hyperoxic environment [112]. Low 2% O_2_ is more physiological to what fetal muscle cells are exposed to *in utero*, allowing “normal” function without being hypoxic [113]. Hypoxia can increase ROS production [114], however culturing fetal muscle cells at 0.5% O_2_ did not display reduced cell growth rates or increase senescent marker expression. The cells may have adapted to growth at low oxygen concentrations, or perhaps 0.5% oxygen was not decreasing intracellular oxygen tension (PO_2_) to a point where oxygen availability limited respiration enough to create hypoxia and oxidative stress [114].

Catalase and SOD1 levels were significantly higher in 20% O_2_ cultured cells compared to the lower oxygen groups, in agreement with previous reports that show hyperoxic conditions increase antioxidant levels [115, 116]. Telomerase inhibition did not significantly alter the levels of these antioxidants, although senescence has been associated with decreased antioxidant capacity [117]. The non-significant increase in Catalase and non-significant decrease in SOD for telomerase inhibited 20% O_2_ cells may together produce decreased amounts of cellular H_2_O_2_ explaining the significant decrease in cellular ROS levels measured after telomerase inhibition. Alternatively, other antioxidants such as mitochondrial SOD2 may show differing effects following telomerase inhibition and could be examined in future studies.

P66Shc is a unique stress adaptor protein, that when phosphorylated at serine-36 translocates into mitochondria to produce and release ROS that can induce apoptosis or senescence [118, 119]. Phospho-p66Shc was increased in fetal muscle cells upon treatment with TI-IX. Telomerase inhibition could regulate p66Shc phosphorylation/activation and mitochondrial ROS production through p53 [120] and/or through ROS activation of p66Shc in a feed forward manner [69, 118]. This links DNA damage to increased ROS production through a p53-p66Shc mechanism that may drive other senescent programs through p66Shc’s regulation of cellular metabolism [70, 121].

Senescent cells have lost the ability to divide, but still carry out cellular functions that bypass programmed cell death [122]. Cell metabolism regulators such as mTOR and S6K have been implicated in senescence [123, 124] so they were investigated along with other metabolic regulators. We observed no difference in mTOR or phospho-mTOR between groups treated with TI-IX. Other phosphorylation sites on mTOR may be altered and could be investigated in future experiments. However, here was a significant difference among oxygen tensions, with increased phospho-mTOR detected at lower oxygen concentrations. Previous studies report that mTOR is down regulated by hypoxia [125], further indicating that our 2% oxygen cultured cells were not hypoxic. mTOR may play a role in elevating telomerase activity of low O_2_ cells, as other studies have shown that increased mTOR and related pathways such as Hsp90 and Akt can increase telomerase levels [126, 127]. Unlike phospho-mTOR, phospho-S6K was significantly up-regulated in 20% O_2_ telomerase inhibited cells. Our results indicate that senescence may be mediated through S6K phosphorylation, and its activation is associated with p66Shc activation as well. Under oxidative stress, p66Shc activation mediates S6K phosphorylation and activates p53, linking mitochondrial function and DNA damage to cellular metabolism [70]. Akt and phospho-Akt were both significantly decreased upon telomerase inhibition, but the phospho-Akt/Akt ratio was not significantly different. There have been numerous conflicting studies for Akt inducing [128] and inhibiting senescence [129, 130]. Akt has been shown to modulate senescence through a downstream kinase, GSK-3 [72]. We observed Phospho-GSK-3 at serine 9 was significantly increased in fetal muscle cells cultured at 20% O_2_ with telomerase inhibition. GSK-3 has been shown to have a role in metabolic dysregulation including increasing cell mass and protein translation along with modulating various cellular organelles [131]. This connects GSK-3 and senescence to S6K and increased protein mass to create large cells that express a metabolic senescent phenotype. In addition to its role in modulating metabolism, active GSK-3 phosphorylates p21 and targets it for ubiquitin-mediated degradation [72]. Our current findings suggest that inactive GSK-3 (serine-9 phosphorylation) would diminish p21 degradation, allowing for p21 levels to accumulate during permanent cell cycle arrest. Together, our findings agree with a number of studies that show a link between nutrient sensing, metabolic dysregulation, oxidative stress and DNA damage that promotes senescence under stressful environments [132–134].

In this study, telomerase inhibition induced premature cellular senescence of high oxygen cultured fetal guinea pig muscle cells. Cells cultured in atmospheric oxygen are in a continued state of oxidative stress, and we argue that these cells can be used for modeling oxidative stress environments *in utero*. Previous studies have shown that hypoxia-reoxygenation or oxidative bursts occur *in utero* [135–137] and modulation of energy substrates can effect fetal growth during development [138]. Our work may help elucidate premature senescence and aging pathways that may be taking place in fetal cells/tissues subjected to adverse *in utero* environments such as those that occur during intrauterine growth restriction (IUGR). Telomerase activity in the placenta is negatively correlated with poor pregnancy outcomes [139]. IUGR has been associated with reduced telomerase activity levels [140] that are associated with decreased placenta size [141], increased oxidative stress markers [142], shortened telomeres or telomere aggregates [55, 59, 62, 143, 144], and increased DNA damage and senescent markers [60, 145, 146]. Independent of their length, telomeres are preferred targets of a steady DNA damage response in aging and stress-induced premature senescence [99]. Our results strengthen the importance of telomerase activity, and if significantly decreased during development may lead to premature senescence and early onset of age-associated diseases in IUGR offspring.

## Materials and Methods

### Fetal guinea pig muscle cell culture

A primary fetal guinea pig myoblast cell line of was derived from the soleus skeletal muscle of a guinea pig euthanized by CO_2_ asphyxiation. All experimental protocols were approved by the University of Western Ontario Animal Care and Veterinary Services and the Canadian Council of Animal Care (animal utilization protocol (AUP) no. 2010-229). Fetal guinea pig muscle cells were cultured in Dulbecco’s Modified Eagle Medium (DMEM) with high glucose and sodium pyruvate. DMEM was supplemented with 10% fetal bovine serum (FBS), 1% L-Glutamine and 1% Penicillin-Streptomycin. Cells were cultured at 37°C under in a humidified 5% CO_2_ atmosphere at either 20% oxygen (O_2_), 2% O_2_ or 0.5% O_2_ tensions. These oxygen tensions estimate the oxygen concentrations found in adult arterial PO_2_ (∼80-100mmHg), fetal arterial PO_2_ (∼15mmHg), and a lower oxygen tension to model a hypoxic arterial PO_2_ (∼3mmHg), respectively [113]. Cells were grown in the same conditions over the entirety of their treatment periods.

### Trypan blue exclusion test of cell viability and determination of population doubling time

Cell viability was manually determined using a hemocytometer with 0.4% trypan blue to stain nonviable cells, with both nonviable and viable cells separately counted. Briefly, an aliquot (∼5 x 10^5^ cells/mL) of cells were centrifuged at 100 x g for 5 min and the supernatant discarded. The cell pellet was resuspended in 1 ml PBS and 1 part 0.4% trypan blue solution and 1 part cell suspension were mixed and incubated for ∼3 min at room temperature. Within 3-5 min cells were counted using a hemocytometer. The following equation was used to calculate cell viability: Viable cells (%) = (total number of viable cells per ml of aliquot / total number of cells per ml of aliquot) x 100. To determine population-doubling times, fetal guinea pig muscle cells were seeded at three different concentrations (25,000, 35,000 and 50,000) in six well dishes and viable cells counted every 24 hours for three days. Cell population doubling times were calculated using the algorithm provided by http://www.doubling-time.com.

### Pharmacological inhibition of telomerase activity

Telomerase Inhibitor (TI)-IX (Calbiochem; EMD Millipore, Billerica, MA) is a cell-permeable, bis-catechol containing m-phenylenediamide compound (MST-312) that inhibits telomerase activity [76, 147]. Fetal guinea pig muscle cells were cultured in the presence or absence (non-treated and DMSO treated (vehicle) controls) of various concentrations (0.1 μM, 0.5 μM, 1.0 μM and 2.0 μM) of TI-IX for 48 hr at 37°C under a 5% CO_2_ atmosphere containing either 20% or 2% O_2_ tension and subsequently examined.

### Real-time quantitative telomeric repeat amplification (RQ-TRAP) Assay

To quantify telomerase activity levels, the TRAPEZE^®^ RT Telomerase Detection Kit (Chemicon; EMD Millipore) was utilized. Cells were collected and lysed to extract total protein and RNA using CHAPS lysis buffer (30 mM Tris-Cl pH 7.5, 150 mM NaCl, 1% CHAPS) containing 100 units/mL RNaseOUT™ (Invitrogen; Life Technologies, Burlington, ON) on ice (20 min) and then centrifuged at 12,000 x g for 20 min at 4°C. Supernatants were assayed for protein content using RC DC Protein Assay Kit II (BioRad, Hercules, CA), and each sample was brought to a final sample concentration of 1.5 μg/μL.

Telomerase activity was assayed using 1.5 μg of protein per reaction, and experiments performed in triplicate. For all experiments, human embryonic kidney cells (HEK293T) were used as a positive control and human dermal fibroblast (HDF) as a negative control. Telomerase, if present in the cell extract, will add telomeric repeat sequences (TTAGGG) sequences to the 3’ end of an oligonucleotide substrate. The number of repeat sequences added by telomerase is quantified using real-time quantitative PCR by measuring the increase in SYBR^®^ green fluorescence upon binding to DNA. The parameters for real-time qPCR were: 30 minutes at 3O°C, 2 minutes at 95°C, followed by 45 cycles of 94°C for 15 seconds, 59°C for 60 seconds, and 45°C for 10 seconds. A total volume of 12 μL was loaded into 394-well plates consisting of 1.0 μL of each sample (1.5 μg) or positive control and 11 μL of master mix. The master mix is comprised of 5x TRAPEZE^®^ RT Reaction Mix, nuclease-free water and Titanium Taq DNA Polymerase (Clontech Laboratories Inc.; Takara Bio Group). Telomerase activity standard curves were generated using four 10-fold serial dilutions (0.02, 0.2, 2.0 and 20 amoles/μL) of the provided telomerase positive control sample (TSR8) with the CHAPS lysis buffer. Calculating the log_10_ of the amoles per well for each reaction and plotting them against the experimental cycle time (Ct) average generated these standard curves that were fitted with a linear trend line and equation. Experimental sample average Ct values were used to extrapolate arbitrary telomerase activity units relative to the TSR8 positive control values in the standard curve.

### Immunofluorescence microscopy

To ensure the primary cell line was indeed muscle cell progenitors, immunocytochemistry was employed using muscle-specific differentiation factors. Fetal guinea pig muscle cells were cultured for 48 hrs in 4-well chamber slides (Nunc; Thermo Scientific, Canada) and then fixed for 10-15 min using 4% paraformaldehyde. Cells were then permeabilized and blocked for 1 hr using a solution of phosphate buffered saline (PBS), 5% Goat Serum (Sigma-Aldrich Co., Canada) and 0.1% Triton X-100. Primary antibodies against muscle specific transcription factors Pax7, Myogenin and MyoD (Santa Cruz Biotechnology Inc., Dallas, TX) were each used in 1:500 dilutions in the blocking solution. The primary antibody incubation was carried out at room temperature for 1 hr, followed by three consecutive washes with 0.1% Triton X-100 in PBS (PBST-X). Appropriate secondary antibodies (anti-mouse or anti-rabbit conjugated with Alexa Fluor-488; Invitrogen) were diluted 1:1000 in blocking solution and Hoescht nuclear. Secondary antibody labelling was completed at room temperature for 1 hr in the dark, followed by three PBST-X washes. Slides were then mounted with cover slips using Anti-Fade Gold Reagent (Invitrogen; Life Technologies) and were visualized and imaged using a Leica DMI6000 inverted microscope (Leica Microsystems Inc., Concord, ON, Canada).

### Telomere-dysfunction Induced Foci (TIFs) detection by immuno-FISH

Telomere-dysfunction induced foci (TIFs) are generated when DNA damage markers co-localize with a telomeric probe to indicate double-stranded DNA damage at the telomeres [148]. In this experiment, control and TI-IX treated cells were grown for 48 hrs on 4-well chamber slides (Nunc), initially washed in PBS and fixed with a 2% paraformaldehyde/2% sucrose solution for 10 min at room temperature. The cells were then permeabilized with 0.5% Nonidet-P40 (NP-40), and washed three times for 5 min. Cells were subsequently incubated for 1 hr at room temperature in a blocking solution consisting of PBS with 5% Goat Serum (Sigma-Aldrich). Following blocking, cells were labeled with a primary antibody against the DNA damage indicator phosphorylated histone ɣH2AX (Phospho-Ser139) (Cell Signaling Technology, Danvers, MA). The primary antibody was diluted 1:2000 in blocking solution and incubated with the cells at room temperature for 1 hr. Cells were washed using PBST-X prior to secondary antibody labeling. Anti-rabbit Alexa-Fluor 488 secondary antibody (Invitrogen) was used at 1:2000 dilution in blocking solution for 1 hr at room temperature in the dark. The primary and secondary antibodies were then post-fixed by treating cells with 4% Paraformaldehyde. The next step utilized a peptide nucleic acid (PNA) probe against telomere sequences in fluorescent *in situ* hybridization (FISH).

The TelC-Cy3 PNA FISH probe (Panagene Inc., Daejeon, Korea) was used in the hybridization buffer with water, 2% Bovine Serum Albumin (BSA), 0.06 ug Yeast tRNA (Invitrogen), 0.6x SSC (diluted from 20x SSC: 3M NaCl, 300mM Trisodium citrate, pH 7.0), and 70% formamide (*see* Supplementary Table 1). Slides were incubated with 120 μL of the PNA-FISH hybridization buffer and were denatured at 85°C for 5 min, and then placed in a humidified chamber in the dark overnight. Twenty-four hours later, slides were washed twice with wash solution I (10mM Tris-HCl, 70% Formamide, 0.1% Tween 20, 0.1% BSA, pH 7.5). This was followed by three washes in solution II (50mM Tris-HCl, 150mM NaCl, 0.1% BSA, 0.1% Tween 20, pH 7.5). Slides were then ethanol dehydrated in consecutive washes with 70%, 85% and 95% ethanol and then stained with Hoescht 33342 nuclear stain for 10 min. Chamber slides were removed and mounted with coverslips and anti-fade gold reagent mounting media. Slides were visualized on a Leica DMI6000 inverted microscope and ORCA-R2 digital camera (C10600, Hammumatsu) in three channels: Leica DAPI, Leica GFP and Leica Texas Red. Images taken from each channel were overlaid to determine the number of cells exhibiting co-localization foci of green and red fluorescence indicating telomeric DNA damage.

### Western blot analysis

Total protein was extracted from fetal guinea pig muscle cells using RIPA buffer (50 mM Tris-HCl, 150mM NaCl, 1mM EDTA, 1% NP-40, 0.25% Na-deoxycholate, pH 7.4) containing protease inhibitors (Roche) and phosphatase inhibitors (5mM sodium fluoride, 5mM sodium beta-glycerophosphate, 1mM sodium orthovanadate, 1mM sodium phyrophosphate). Cells were mechanically homogenized with a pestle and incubated on ice for 10 min. Extractions were centrifuged at 4°C at 12,500 x g for 10 min and the supernatant was collected and frozen at −20°C until further analysis.

Nuclear fractions were extracted from cells using Buffer A with protease and phosphatase inhibitors. Cells were mechanically homogenized with a pestle and incubated on ice for 10 min. Extractions were centrifuged at 4°C at 300 x g for 10 min. Cell pellets were discarded and the supernatant was collected and centrifuged again at 1500 x g for 10 min. Supernatant was discarded and the nuclear fraction was resuspended in RIPA buffer and sonicated to break up DNA in nuclear fraction. Samples were frozen at −20°C until further analysis.

Samples from protein extractions were quantified using the RC DC Protein Assay Kit II (BioRad) by using a Spectramax spectrophotometer (Molecular Devices, Sunnyvale, CA). The samples were prepared with NuPage LDS Sample Buffer and NuPage Sample Reducing Buffer (Invitrogen). Twenty μg of total protein extracts and 10 μg of nuclear extractions were loaded on 4-12% Bis-Tris gels (Invitrogen) in SDS-MES running buffer. Gel electrophoresis was run at 180V for 45-60 min until proteins were separated. Proteins were transferred from Bis-Tris gels to polyvinylidene fluoride (PVDF) membranes for 2 hours using a wet transfer system (BioRad). Following transfer, membranes were blocked in either 5% skim milk or 5% BSA in Tris-buffered saline with Tween 20 (TBST) (50 mM Tris-HCl, 150 mM NaCl, 0.1% Tween 20, pH 7.6) for 1 hr. Following blocking, membranes were incubated overnight at 4°C on a rotator in primary antibodies of 1:1000 dilutions (*see* Supplementary Table 2). Housekeeping proteins: β-actin (total protein extractions) and histone H3 (nuclear protein extractions) were also used at higher dilutions of 1:20000 and 1:5000, respectively. Membranes were washed with TBST prior to incubation for 2 hrs at room temperature in the appropriate secondary antibody (anti-rabbit or anti-mouse; Cell Signaling) at 1:2000 dilutions. Membranes were washed with TBST before detection using SuperSignal West Pico Chemiluminescent Substrate (ThermoScientific). Protein bands were visualized using the VersaDoc Imaging System (BioRad) and quantified using Image Lab Software (BioRad).

### Annexin V and propidium iodide staining of apoptotic cells

To determine if telomerase inhibition triggered apoptosis, an apoptosis kit was used containing Annexin-V, a cell surface marker for apoptosis, and a dead cell nuclear marker propidium iodide (PI) (Invitrogen; Life Technologies). Twenty-four hrs prior to analysis cells were treated with 500 μM hydrogen peroxide (H_2_O_2_) as a positive control treatment. Cells were resuspended in annexin-binding buffer to a concentration of 1.0 x 10^6^ cells/ml in 100 μl. Alexa-fluor 488 annexin-V was added at a dilution of 1:20 and PI was added to a concentration of 0.01 μg/mL and incubated at room temperature for 15 min in the dark. Following incubation, 400 μL annexin binding buffer was mixed with each sample and kept on ice until analysis by the Accuri C6 flow cytometer (see details below). Alexa Fluor 488 annexin-V fluorescence excitation/emission maxima were measured at 488/530-575 nm, and PI was measured at 535/617 nm.

### Mitochondrial membrane potential and intracellular ROS production

To assess mitochondrial membrane potential and reactive oxygen species (ROS) production multiple markers were used including MitoSOX™ (Invitrogen), Mitotracker® Red CMXRos (Invitrogen) and carboxy-H2DCFDA (DCF) (Invitrogen). All fluorescent markers were resuspended in dimethyl sulfoxide (DMSO) and incubated with cells for 15 min followed by trypsinization and resuspension in PBS. Cells were analyzed at a concentration of 1.0 x 10^6^ cells/ml using the Accuri C6 flow cytometer (see details below). MitoSOX and DCF were used at a final concentration of 5 μM while Mitotracker CMXRos was used at a final concentration of 0.2 μM. Fluorescent emission/excitations were as follows: 510/580 nm for MitoSOX, 579/599 nm for Mitotracker Red CMXRos and 492-495 nm/517-527 nm for DCF.

### Flow cytometry analysis

Following fluorescence staining protocols, resuspended cells were examined using the Accuri C6 Flow Cytometer (BD Biosciences; Mississauga, Canada). A total of 50,000 events were collected from each sample and these samples were gated to include viable cells as determined by size using forward scatter (FSC) and side scatter (SSC) (S3 Fig). The gated samples were then applied to fluorescent emissions, and the mean fluorescence of the cells in the gated samples were measured and statistically analyzed.

### Statistical Analysis

All data presented is the mean ± standard error of mean (SEM). Data and statistical analysis was conducted using Graph Pad Prism. A one-Way ANOVA with Tukey’s post hoc test was used to compare telomerase inhibition in samples in the same oxygen concentration. A two-Way ANOVA with Bonferroni post hoc test was used to compare telomerase inhibition across different oxygen concentrations. Statistically significant data was indicated if the P-value was 0.05 or less (p<0.05).

## Acknowledgments

The authors would like to thank Lin Zhao for his technical support. This research was supported by the Canadian Institutes of Health Research (CIHR), a Discovery grant to DHB from the Natural Sciences and Engineering Research Council of Canada (NSERC) and a Lawson Research Institute Internal Research Fund to DHB and TRHR. A QEII Graduate Scholarship in Science and Technology (QEIIGSST) funded SEH.

## Supporting information

**S1 Fig. Muscle differentiation markers MyoD, Myogenin and Pax7 indicate muscle lineage cells from fetal guinea pig soleus muscle.** Following 48 hours of incubation at 20% oxygen, fetal guinea pig muscle cells were fixed on 4-well chamber slides and incubated with primary antibodies: MyoD, Myogenin, and PAX7 and subsequently detected using the appropriate Alexa Fluor-488 conjugated secondary antibodies. Fixed cells were counterstained with Hoescht 33342 and the corresponding images are represented in the right column. The scale bar in the bottom right corner represents 50 μM.

**S2 Fig. DCF fluorescence decreases with lower oxygen concentration and telomerase inhibition.** DCF was incubated on cells for 15 mins following a 48-hr treatment with or without Telomerase Inhibitor IX at 20% and 2% oxygen. Cells were then visualized by fluorescence microscopy as shown in (A) or analyzed with the Accuri C6 flow cytometer. Mean fluorescence was determined and represented graphically in (B). Results were analyzed with a two-way ANOVA, n=3, and significant differences are indicated by (*) and (***), representing p<0.05, and p<0.001, respectively. Scale bars represent 50 µM.

**S3 Fig. Flow cytometry analysis using MitoSox with the Accuri C6.** Fetal guinea pig muscle cells were stained for 15 min with MitoSox and resuspended in PBS prior to running through the Accuri C6 flow cytometer. (A) 50,000 events were collected and were plotted by size using forward scatter (FSC; x-axis) and side scatter (SSC; y-axis). Based on size, cells could be gated and cellular debris can be removed. (B) Example of the red fluorescence channel FL2 without gating. (C) The same fluorescence with gating applied showing fluorescence only exhibited by viable cells included. The mean fluorescence was measured and used in statistical analyses.

